# Short-term synaptic dynamics in the ventrolateral and dorsomedial periaqueductal gray

**DOI:** 10.1101/2025.08.12.669947

**Authors:** A.V. Castro Romero, K.A. Tyner, J.N. Carroll, C.E. Vaaga

## Abstract

The ability to assess and rapidly respond to predator threats in the environment is necessary for survival and requires dedicated neural circuits for threat detection, sensorimotor integration, and execution of ethologically appropriate behavioral responses. Although numerous brain circuits are involved in these processes, the midbrain periaqueductal gray (PAG) serves as an important central hub to generate ethologically appropriate passive and/or active defensive behaviors. Despite its central role in generating defensive behaviors, little is known about the intrinsic and synaptic properties of neurons across columns in the PAG. To address this knowledge gap, we made whole-cell voltage- and current-clamp recordings from unlabeled neurons in the vl- and dmPAG of mice. Consistent with *in vivo* work, our data highlights the relative importance of synaptic inhibition in both columns. Further, our results suggest that neurons in both the vl- and dmPAG prioritize frequency-invariant coding strategies, showing remarkably stable paired pulse ratios across interstimulus intervals. Despite this common theme, the underlying mechanism each column utilizes to achieve such frequency invariant coding is distinct, reflecting important differences in synaptic processing across columns. More specifically, while the vlPAG is relatively resistant to phasic short-term depression across stimulation frequencies, neurons in the dmPAG show a pronounced buildup of tonic/slow current during high frequency stimulation trains, which counteracts short-term depression of the phasic current amplitude observed during high frequency stimulation trains. This prolonged tonic current observed in the dmPAG prolongs the period of spike elevation, suggesting that high frequency stimulation may drive sustained activity in the dmPAG. Together, these results provide fundamental information of synaptic integration and network properties across columns in the PAG, which ultimately support their distinct roles in threat processing.

## Introduction

The midbrain periaqueductal gray (PAG) plays an important role in coordinating and generating defensive behavioral reactions to innate and conditioned threats mediated in part through downstream projections to brainstem circuits (Fanselow, 1991; Tovote *et al*., 2016; Deng *et al*., 2016; Assareh *et al*., 2016; Vaaga *et al*., 2020). Anatomically, the PAG can be subdivided into multiple rostro-caudal columns, each of which has classically been associated with driving passive (i.e. freezing; mediated by the vlPAG) or active (i.e. flight; mediated by the dmPAG) coping behaviors in response to threat (Bandler *et al*., 1985; Bandler & Carrive, 1988; Bandler & Shipley, 1994; Tovote *et al*., 2016; Deng *et al*., 2016; Vaaga *et al*., 2020; La-Vu *et al*., 2022). However, recent *in vivo* functional data has begun to challenge the view that PAG circuits simply act as relays, instead identifying more nuanced roles in sensorimotor integration and threat processing (Evans *et al*., 2018; Wright & McDannald, 2019; Walker *et al*., 2019; Reis *et al*., 2021; Strickland & McDannald, 2022), perhaps in addition to more traditional roles in driving behavioral fear responses. For example, *in vivo* data suggests that circuits in the dorsomedial PAG may perform synaptic thresholding computations during approach behaviors, which may play a critical role in initiating fear responses regardless of the coping strategy (Evans *et al*., 2018). Furthermore, in more complex fear tasks, the majority of neurons in the vlPAG encode threat probability over behavioral activity (Wright & McDannald, 2019), suggesting a role in threat integration. These functional studies point to an underappreciated role of PAG circuitry in sensorimotor integration and threat assessment, raising important, yet unresolved, questions of how each columns processes information.

Numerous pre- and postsynaptic factors can impact information processing dynamics in neural circuits. For example, activity dependent changes in synaptic strength over short time scales (ms – min) can have a profound influence on the synaptic computations performed within a neural circuit and play a critical role in determining how neurons and circuits process information (Abbott & Regehr, 2004). Specific profiles of short-term synaptic dynamics, which include pre- and postsynaptic factors (Trussell & Fischbach, 1989; Dittman & Regehr, 1998; Rozov & Burnashev, 1999; Dittman *et al*., 2000; Geiger & Jonas, 2000; Zucker & Regehr, 2002; Chen *et al*., 2002; Abbott & Regehr, 2004), provide neural circuits with an enhanced capacity for information processing. For example, presynaptic release probability is one of the primary drivers of short-term synaptic plasticity and can result in differential synaptic filtering and ultimately distinct circuit function (Markram *et al*., 1998b, 1998a; Dittman *et al*., 2000; Silberberg *et al*., 2004; Abbott & Regehr, 2004). More specifically, synapses are often categorized into one of two groups: facilitating or depressing. Synapses with a high initial release probability tend to undergo paired pulse depression (acting as a low-pass filter) whereas synapses with a low release probability undergo facilitation (acting as a high-pass filter). Such characterizations of synapse function, although helpful, do not fully capture the breadth of synaptic dynamics within neural circuits, as synaptic transmission can also engage band-pass filters through intermediate release probabilities or engage in frequency invariant synaptic transmission (Dittman *et al*., 2000; Armstrong-Gold & Rieke, 2003; Telgkamp *et al*., 2004; Abbott & Regehr, 2004; Turecek *et al*., 2016, 2017; Mondal *et al*., 2022). Furthermore, understanding additional features of circuit function, including biophysical specializations, intrinsic excitability, synaptic excitation/inhibition (E/I) ratios, and relative ability to engage in synaptic plasticity are necessary to provide a framework with which to interpret the functional impact of synaptic input. Together these factors point to the importance of elucidating the synaptic dynamics across and within the periaqueductal gray columns to understand how each column processes information and ultimately contributes to the generation of ethologically relevant fear responses.

Here we characterize and compare the intrinsic properties and short-term synaptic dynamics of neurons in the ventrolateral (vl-) and dorsomedial (dm-) PAG. We demonstrate that in both circuits inhibitory strength outweighs excitation, reinforcing the importance of inhibitory tone within the PAG (Tovote *et al*., 2016). Further, using paired pulse stimulation paradigms we show that both inhibitory and excitatory efficacy are remarkably stable across interstimulus intervals, without appreciable change in the paired pulse ratio across intervals, suggesting frequency invariant synaptic coding. Interestingly, despite this commonality, we find significant differences in the underlying synaptic mechanisms utilized by each column to achieve such frequency invariance. Together, our data identify synaptic coding strategies which are conserved across columns, but further identify latent differences in the underlying synaptic mechanisms, which may have important implications for column specific threat processing.

## Methods

### Animal Care and IACUC Approval

All experimental animal procedures were conducted in accordance with institutional guidelines regarding the ethical use of animals. Experimental methods were approved by the Colorado State University Institutional Animal Care and Use Committee (protocol 3820 and 3836, CEV). Wildtype mice (genetic background: C57Bl/6J; age >4 weeks) used in the study were either purchased directly from Jackson Laboratories (strain: 000664) or bred in-house. Experiments were balanced across sex and data from both sexes were pooled for analysis.

### Slice Preparation

Acute coronal slices of the PAG were collected as described previously (Vaaga *et al*., 2020). Briefly, animals were deeply anesthetized using isoflurane until the absence of spinal reflexes. To improve tissue health, mice were transcardially perfused with ∼10 mL of cold (4°C), oxygenated modified artificial cerebral spinal fluid (ACSF) which contained (in mM): 83 NaCl, 2.5 KCl, 1 NaH_2_PO_4_, 26.2 NaHCO_3_, 22 glucose, 72 sucrose, 0.5 CaCl_2_, 3.3 MgCl_2_ (pH 7.2, 290-300 mOsm). After perfusion, the brain was removed and the PAG blocked in the coronal plane at the levels of the cerebellum and optic chiasm. The tissue block was then embedded in 2-4% agarose before slices (300 μm) were made using a compresstome (Precisonary Instruments). Slices were then removed from the agarose embedding and transferred to a warm (37°C) recovery chamber with normal, oxygenated ACSF for at least 30 minutes. The ACSF contained (in mM): 123 NaCl, 3.5 KCl, 1.25 NaH_2_PO_4_, 26 NaHCO_3_, 1 MgCl_2_, 1.5 CaCl_2_, 10 glucose (pH 7.2, 290-300 mOsm). After 30 minutes, slices were maintained in normal ACSF at room temperature (22-23°C) until use.

### Electrophysiological Recordings

Current-clamp and voltage-clamp recordings were made from neurons in the vl- and dmPAG under DIC optics using an Olympus BX51WI microscope. For current clamp recordings, a K-gluconate based internal solution was used, which contained (in mM): 130 K-gluconate, 2 Na-gluconate, 6 NaCl, 2 MgCl_2_, 0.1 CaCl_2_, 1 EGTA, 4 MgATP, 0.3 trisGTP, 14 tris-creatine phosphate, 10 sucrose, 10 HEPES (pH adjusted to 7.35 with KOH; 287 mOsm). For voltage clamp recordings a Cs-MeSO_3_ internal solution was used, which contained (in mM): 120 CsMeSO_3_, 3 NaCl, 2 MgCl_2_, 1 EGTA, 10 HEPES, 4 MgATP, 0.3 trisGTP, 14 tris-creatine phosphate, 12 sucrose and 1.2 QX314 (pH adjusted to 7.32 with CsOH, 287 mOsm).

Borosilicate patch-pipettes were pulled to a resistance of 4-6 MΩ using a Narishige PC-100 two-stage vertical puller. During recordings, the temperature of recording chamber was maintained at 34-36°C by a Warner TC-324B controller. Data were acquired using a Sutter Instruments Integrated Patch Amplifier (IPA). Data were acquired at 20 kHz and filtered at 10 KHz. The liquid junction potential (10 mV) was corrected during recordings. During voltage clamp recordings, access resistance was monitored by a 10 mV hyperpolarizing step, and recordings with a >20% change in access resistance were discarded from analysis. For current clamp recordings, access resistance was compared before and after the recording, and cells with a >20% change in access resistance were discarded from further analysis. Additionally, cells with high holding current (generally above ∼300 pA), at any point in the recording, were discarded from analysis.

Spontaneous action potential recordings were acquired in current clamp configuration with 10 pA of injected current. To construct FI curves, cells were injected with 500 ms steps of current (− 100 pA to 200 pA). In voltage clamp recordings, synaptic currents were evoked using a parallel bipolar electrode (FHC instruments) inserted into the tissue lateral to the recording site (200 – 300 μm) in either the vlPAG or dmPAG. Electrical stimulation (0.1 ms, 0.7-3.2 mA) of either polarity was used, and the intensity of the stimulus was adjusted to produce reliable evoked currents. Unless otherwise noted, EPSCs and IPSCs were isolated using reversal potential (V_cmd_ = 0 mV to isolate IPSCs, V_cmd_ = −70 mV to isolate EPSCs). Synaptic currents were evoked with an interstimulus interval of >5 seconds and between 5-20 sweeps were collected in each condition. For trains of stimuli, stimulation frequency varied between 10 Hz and 100 Hz (10 pulses per sweep). All drugs were acquired from Tocris Biosciences and were bath applied at the following concentrations: 10 μM DNQX (6,7-dinitroquinoxaline-2,3-dione), 10 μM CPP (3- ((*R*)-2-Carboxypiperazin-4-yl)-propyl-1-phosphonic acid), 10 μM SR95531 (6-Imino-3-(4-methoxyphenyl)-1(6H)-pyridazinebutanoic acid hydrobromide).

### Data Analysis

Electrophysiological data was analyzed using IgorPro 9 (Wavemetrics, Sutter Instruments) in combination with the SutterPatch software package. For action potential analysis, we applied an amplitude threshold (0 mV) and measured the following properties: half-width, threshold, maximum dV/dt, minimum dV/dt, and peak amplitude. FI curves were constructed by measuring the number of action potentials at each current step (500 ms) and converting to spike frequency.

For voltage clamp recordings, all data was baseline-subtracted and sweeps were averaged together prior to analysis. Stimulus artifacts were manually removed for visual clarity. To measure synaptic currents, the peak amplitude was measured relative either to the sweep baseline (total current) or to the current immediately prior to the evoked response (phasic current). Spontaneous current records were additionally filtered at 3 kHz and individual events were detected using the SutterPatch synaptic event detection (excitation template: rise time: 500 μs; decay time: 2 ms; inhibition template: rise time: 500 μs; decay time: 5 ms) with threshold of 2x the standard deviation of the noise. Events were then manually reviewed and accepted/rejected on the basis of amplitude and kinetic profile. To calculate the AMPA/NMDA ratio, in each cell, we measured the peak synaptic current recorded at −70 mV (AMPA receptor component) and the current amplitude 40 ms after the peak at +40 mV (NMDA receptor component)

To quantify the contribution of phasic and tonic current components to excitatory responses across stimulation trains we separately measured the phasic current and total current for each EPSC. The phasic current was measured by calculating the peak EPSC on each stimulus relative to the baseline current immediately prior to the stimulus artifact. Conversely, the total current was measured by calculating the peak EPSC on each stimulus relative to the baseline current prior to the stimulus train. By subtracting the phasic current from the total current, we calculated the tonic current (or envelope current) on which phasic EPSCs were generated. Additionally, to quantify the magnitude of the tonic current, we measured the tonic current just prior to the last stimulus in the train and normalized it to the amplitude of the phasic current of the first stimulus.

### Statistical Treatment

Data are reported in the text as mean ± standard deviation and graphed using violin plots with individual data points (cells) plotted and median (solid line) and inter-quartile range (dashed line). To plot data across stimuli, we plotted the mean ± 95% confidence interval. We report both the number of cells (n) as the biological replicate but also include the number of animals (N) from which the data were obtained. The number of recorded cells (at least 10 cells per experiment) reflects the statistical power (α = 0.05, β = 0.8) necessary to detect a 20% effect size using paired statistics based on historical data of synaptic events. Statistical tests were performed in Prism 10 and included (as appropriate): Student’s paired and unpaired t-tests, one-sample t-tests, one-way and two-way ANOVAS (including ordinary and repeated measures). If necessary, we used a Šídák’s post-hoc test to control for multiple comparisons.

## Results

### Comparing the intrinsic firing properties of cells within the vl- and dmPAG

To fully appreciate information processing and synaptic integration within a neural circuit requires understanding the underlying intrinsic properties of the circuit in question. Therefore, we first sought to compare the intrinsic electrophysiological properties of neurons in the vl- and dmPAG (vlPAG: n = 13, N = 6; dmPAG: n = 12, N = 4); more specifically characterizing the intrinsic membrane and firing properties using whole-cell current-clamp recordings (**Table 1**). Across columns, we observed no significant differences in either the membrane resistance (vlPAG: 633.28±257.33 MΩ; dmPAG: 617.80±323.42 MΩ; unpaired t-test: p = 0.89) or cell capacitance (vlPAG: 29.15±12.36 pF; dmPAG: 23.25±6.0 pF; unpaired t-test: p = 0.15). Neurons in both columns were spontaneously active at moderate firing rates (**Figures 1A-B, G**; vlPAG: 15.17±14.63 Hz; dmPAG: 10.78±7.60 Hz); however, there were no differences in the firing rate across columns (**Figure 1G**; unpaired t-test: p = 0.36). Furthermore, we observed no difference in the resting membrane potential, although there was a trend towards more depolarized resting membrane potential in the vlPAG (**Figure 1H**; vlPAG: −53.59±6.89 mV; dmPAG: −58.16±4.30 mV; unpaired t-test: 0.061).

**Figure 1:**
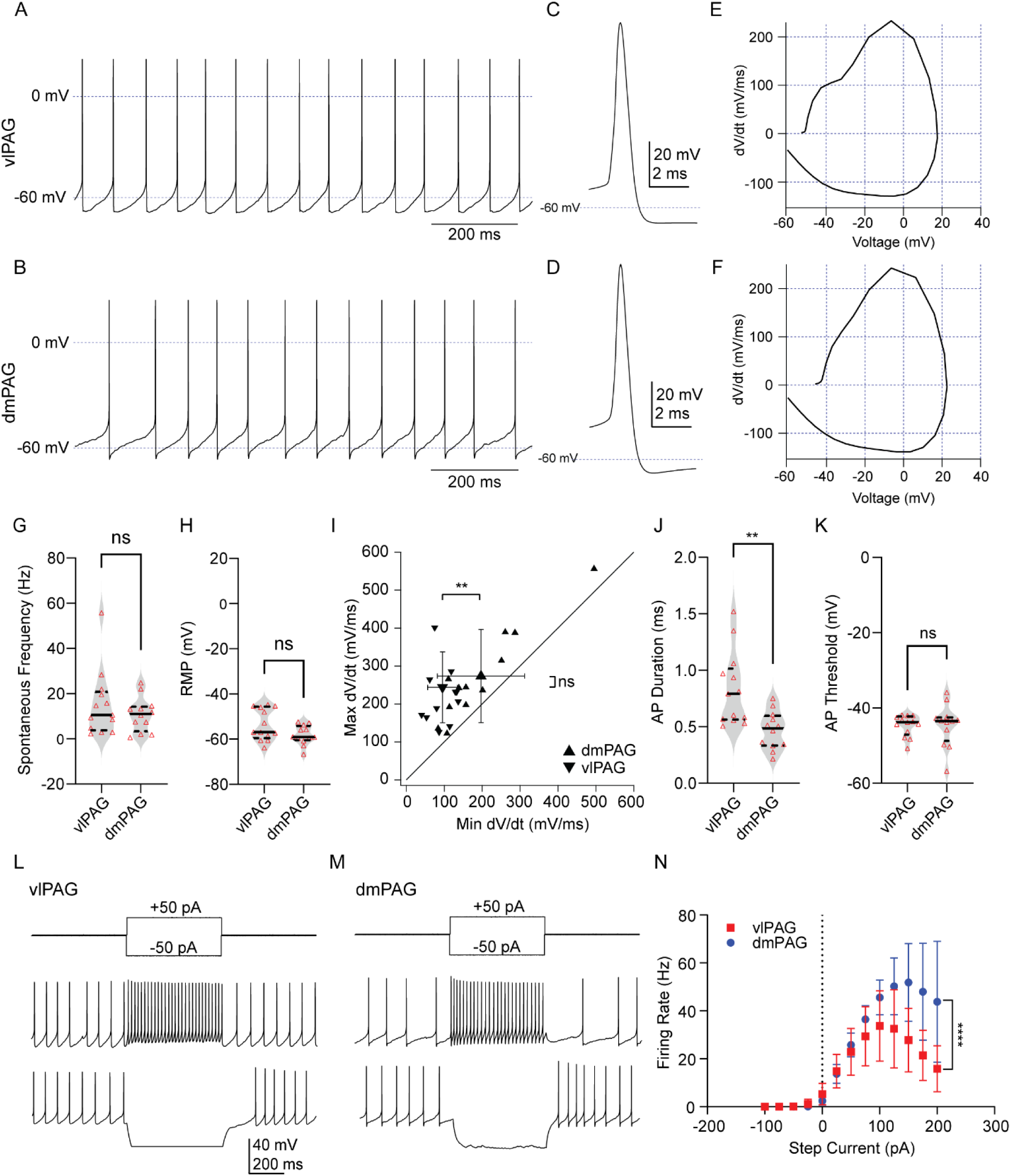
Comparison of intrinsic firing properties of cells within the vl- and dmPAG. **(A, B)** Spontaneous action potentials from unlabeled neurons in the vlPAG (A) and dmPAG (B). **(C, D)** Representative averaged action potential recorded a neuron in the vlPAG (C) and dmPAG (D). **(E, F)** Representative phase plane plot of action potentials recorded within the vlPAG (E) and dmPAG (F). **(G)** Violin plot showing the distribution of spontaneous action potential firing frequency among cells in the vl- and dmPAG. **(H)** Resting membrane potential (RMP) comparison between cells in the vlPAG and the dmPAG. **(I)** Scatter plot of the maximum dV/dt vs. minimum dV/dt recorded from neurons across columns. *Small triangles,* individual cells; *Large triangles,* mean ± SD; vl- (upward facing triangles) /dm- (downward facing triangles) cells. **(J)** Comparison of action potential duration between cells within the vl- and dmPAG. **(K)** Violin plot comparing the action potential threshold between cells within the vl- and dmPAG. **(L, M)** Action potentials evoked by step current injection in the vlPAG (L) and dmPAG (M). **(N)** Averaged evoked firing rate (mean ± SD) in response to current injection in neurons from the vlPAG (red squares) and dmPAG (blue circles).

**Table 1:** Intrinsic properties across columns. Summary of intrinsic membrane properties and action potential properties across neurons in the vlPAG and dmPAG. Reported p-value represents unpaired t-test across columns.

| Parameter | All Cells<br>(n=25) | vIPAG (n=13) | dmPAG (n=12) | p value |
| --- | --- | --- | --- | --- |
| **Intrinsic Properties** |  |  |  |  |
| Membrane Resistance (MΩ) | 625.9 ± 284.8 | 633.56 ± 257.28 | 617.76 ± 323.42 | 0.894 |
| Capacitance (pF) | 26.35 ± 10.06 | 29.31 ± 12 | 23.14 ± 5.77 | 0.122 |
| Resting Membrane Potential | -55.78 ± 6.13 | -53.59 ± 6.89 | -58.16 ± 4.28 | 0.060 |
| **Action Potential Properties** |  |  |  |  |
| Spontaneous Rate (spikes/s) | 13.06 ± 11.76 | 15.17 ± 14.63 | 10.78 ± 7.60 | 0.362 |
| Afterhyperpolarization (mV) | -65.07 ± 4.84 | -63.45 ± 4.55 | -66.84 ± 4.69 | 0.079 |
| Maximum dV/dt (V/s) | 258.01 ±<br>107.22 | 243.7 ± 93.49 | 273.5 ± 122.7 | 0.503 |
| Minimum dV/dt (V/s) | -144.19 ±<br>97.53 | -95.43 ± 39.0 | -197.0 ± 115.1 | 0.006 |
| AP duration (ms) | 0.65 ± 0.31 | 0.83 ± .32 | 0.46 ± 0.16 | 0.002 |
| Amplitude (mV) | 21.43 ± 9.13 | 22.73 ± 10.2 | 20.03 ± 8 | 0.468 |
| Threshold (mV) | -44.64 ± 4.27 | -44.68 ± 2.79 | -44.6 ± 5.6 | 0.96 |

We further characterized the action potential waveform of neurons in both the vl- and dmPAG (**Figure 1C-D**) to ascertain potential differences in intrinsic biophysical properties across columns. Action potential phase-plane analysis, which measures (among other features) the rate of depolarization and repolarization during an action potential, revealed significant differences in the action potential waveform across neurons in the vl- and dmPAG columns (**Figure 1E-F**). More specifically, although the maximal dV/dt rate (a measure of the depolarization rate) was similar across columns (**Figure 1I**; vlPAG: 243.70±93.49 mV/ms, dmPAG: 273.50±122.7 mV/ms, unpaired t-test: p = 0.50), neurons in the dmPAG had a significantly larger minimum dV/dt rate (**Figure 1I**; vlPAG-95.43±39.00 mV/ms, dmPAG: - 197.0±114.1 mV/ms, unpaired t-test: p = 0.0063), indicative of increased K^+^ conductance upon action potential repolarization (Bean, 2007). Consistent with the larger minimum dV/dt, action potentials from dmPAG neurons were also significantly shorter in duration (**Figure 1J**; vlPAG: 0.83±0.33 ms, dmPAG: 0.47±0.17, unpaired t-test: p = 0.0023). Finally, we observed no significant difference in the action potential threshold across columns (**Figure 1K**; vlPAG: - 44.68±2.79 mV, dmPAG: −44.60±5.61 mV; unpaired t-test: p = 0.96). Taken together, these results suggest that while neurons across both columns have comparable intrinsic properties and spontaneous firing rates, there are subtle differences in the action potential waveform across columns, which may be mediated by distinct complements of voltage gated potassium channels.

As an additional measure of neuronal excitability, we also examined the response to current injection (−100 pA to +200 pA, 25 pA increments; **Figure 1L, M**), allowing us to construct an FI curve. Our analysis revealed that neurons in the dmPAG showed significantly enhanced excitability, reflected in the higher maximal firing rate (**Figure 1N**; two-way ANOVA: current step x PAG column interaction: p <0.0001). In both columns, numerous cells (8/13 vlPAG neurons; 7/12 dmPAG neurons) exhibited depolarization block, resulting in a reduction in the number of action potentials produced in response to our largest current steps. To circumvent this complication, we directly compared the maximal firing rate prior to cells entering depolarization block, which was significantly higher in neurons in the dmPAG (vlPAG: 41.38±24.10 Hz, dmPAG: 67.17±22.63 Hz; unpaired t-test: p = 0.011), consistent with their elevated intrinsic excitability. Together these data suggest that the intrinsic excitability of dmPAG neurons is significantly higher than neurons in the vlPAG, perhaps supporting distinct column-specific functional integration.

### Characterization of local and extrinsic synaptic transmission across vl- and dmPAG

To understand whether and how synaptic information is differentially integrated within distinct PAG columns requires, first, characterizing the features of basal synaptic transmission in each column. To address this question, we first attempted to characterize local circuit dynamics by measuring and quantifying the strength and frequency of spontaneous excitatory and inhibitory synaptic currents. Such measurements likely capture local circuit dynamics, as they are preferentially influenced by local, intact synaptic connections preserved within the slice. To quantify the strength and frequency of the local synaptic network, we recorded both spontaneous excitatory and inhibitory postsynaptic currents in individual cells, using the holding voltage protocols to isolate synaptic currents (**Figure 2A, D**). In both columns, the amplitude of spontaneous inhibitory currents were significantly larger than excitatory currents (vlPAG: sEPSC: 21.83±7.10 pA; sIPSC: 34.97±10.11 pA; paired t-test: p = 0.0021, n = 14, N = 2; dmPAG:; sEPSC: 16.16±3.94 pA; sIPSC: 28.08±7.81 pA; paired t-test: p = 0.0001, n = 12, N = 2; **Figure 2B, E**), likely reflecting a higher unitary conductance of inhibitory synaptic currents. Interestingly, there was no significant difference in the spontaneous frequency of inhibitory and excitatory events in either the vlPAG (sEPSC: 2.07±1.14 Hz; sIPSC: 2.98±2.64 Hz; paired t-test: p = 0.2, n = 14, N = 2) or dmPAG (sEPSC: 4.00±2.72 Hz, sIPSC: 3.62±4.45 Hz; paired t-test: p = 0.68, n = 12, N = 2; **Figure 2C, F**). Furthermore, we observed no significant differences in the inhibitory or excitatory spontaneous frequency across columns (two-way ANOVA: main effect of E/IPSC: p = 0.74; main effect of column: p = 0.12; current x column interaction: 0.43), suggesting that the overall local network connectivity is likely equivalent across columns. In summary, these findings suggest that the strength of local inhibition is significantly larger than excitation in both columns. This difference is likely to reflect a larger inhibitory unitary conductance rather than a difference in synaptic connectivity, although differences in spontaneous firing rates across neurochemically diverse presynaptic neurons may compensate for any differences in structural connectivity.

**Figure 2.**
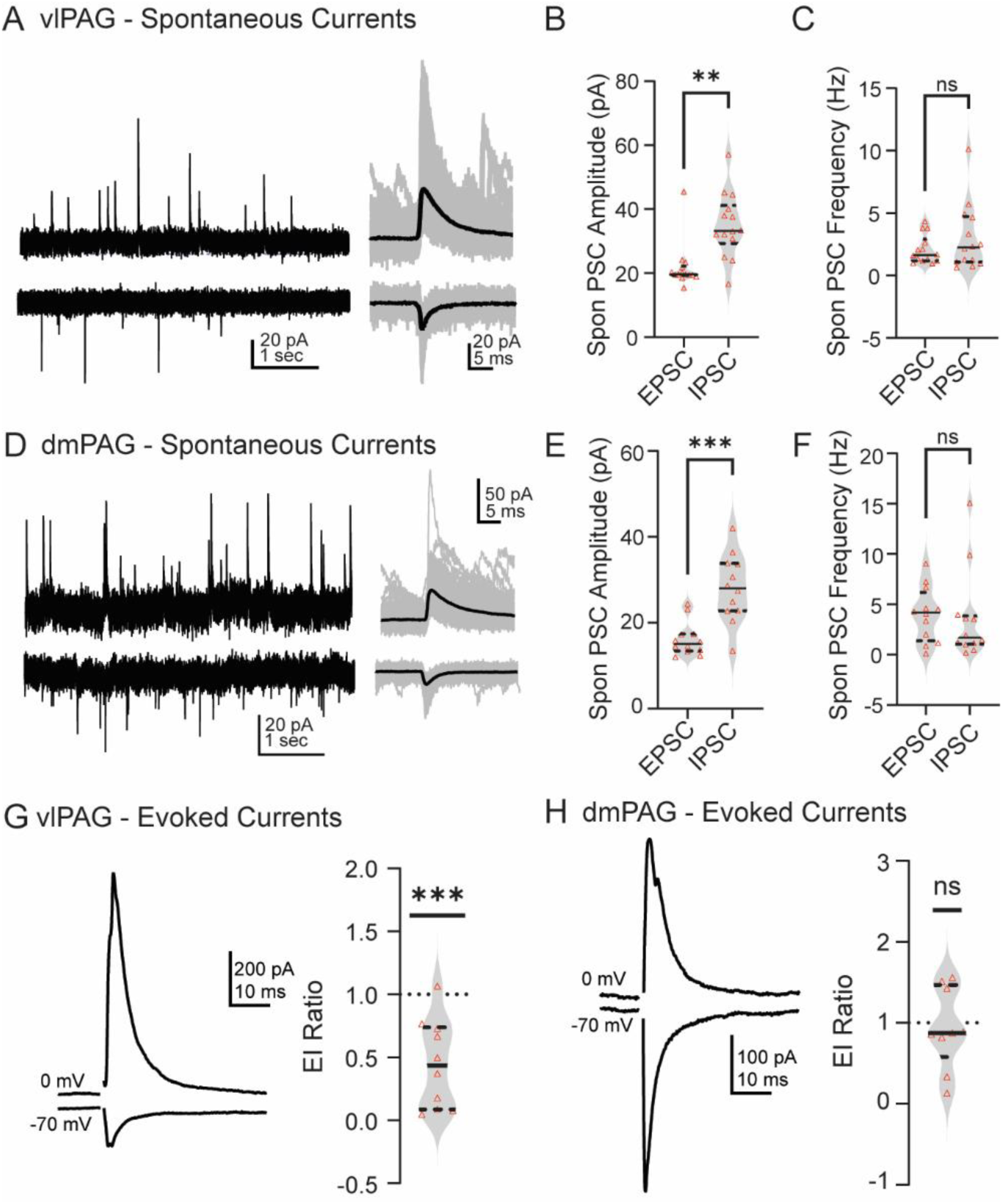
Inhibition outweighs excitation in vl- and dmPAG columns. **(A)** Representative traces of spontaneous excitatory and inhibitory currents (left) in vlPAG recorded at −70 mV and 0 mV, respectively. Isolated individual events (right; gray) with averaged synaptic current overlaid (black). **(B, C)** Violin plots of EPSC and IPSC amplitude (B) and frequency (C) within the vlPAG (solid black line: median, dotted black line interquartile range). **(D)** Representative traces of spontaneous excitatory and inhibitory currents (left) in dmPAG recorded at −70 mV and 0 mV, respectively. Isolated individual events (right; gray) with averaged synaptic current overlaid (black). **(E, F)** Violin plots of EPSC and IPSC amplitude (E) and frequency (F) in the dmPAG. **(G, H)** Representative evoked currents (left) and corresponding distribution of EI ratios (right) in vlPAG (G) and dmPAG (H).

Although spontaneous excitatory and inhibitory currents are well-suited to capture local circuit dynamics, extrinsic inputs are likely underestimated. Therefore, to examine the integration of extrinsic inputs to the PAG, we compared the relative strength of excitation and inhibition following electrical stimulation, which is likely to stimulate both local circuits as well as extrinsic inputs. Specifically, we measured the evoked excitatory (eEPSC) and inhibitory (eIPSC) current within a single cell, allowing us to construct an E/I ratio for each cell. In the vlPAG, eEPSCs had an average amplitude of 80.4±53.0 pA (n = 10, N = 5 whereas eIPSCs had an average amplitude of 294.4±268.8 pA (n = 10, N = 5 **Figure 2G**, left). The stronger inhibition resulted in an E/I ratio significantly smaller than 1 (E/I ratio: 0.45 ±0.35, one-sample t-test: p-0.0008; **Figure 2G**, right). Conversely, in the dmPAG excitation and inhibition were equivalent in amplitude (eEPSC: 405.4±427.0 pA; eIPSC: 359.2±227.0 pA; **Figure 2H**, left; n = 9, N = 3). This equivalency in evoked current amplitude resulted in an E/I ratio that was not significantly different from 1 (E/I ratio: 0.93±0.50, one-sample t-test: p = 0.70; **Figure 2H**, right). Taken together, these data suggest that while local circuit arrangement may be similar across columns, there are likely significant differences in extrinsic circuit transmission across the vl- and dmPAG, consistent with differential extrinsic connectivity at the anatomical level (Beitz, 1982; Silva & McNaughton, 2019). The significantly larger eIPSC amplitude recorded in the vlPAG likely reflects significant extrinsic inhibitory input to the vlPAG, consistent with long-range inhibitory input from the central amygdala (Tovote *et al*., 2016).

### AMPA/NMDA ratios across vl- and dmPAG columns

In addition to quantifying local and extrinsic synaptic strength, another key feature of synaptic integration within neural circuits is the lability of their synaptic plasticity. As a first pass metric of synaptic plasticity capacity across columns, we assessed the AMPA/NMDA ratio across columns, specifically with the goal of quantifying the strength of NMDA receptor-mediated synaptic currents, a key component driving synaptic plasticity in many circuits (Malenka, 1994; Lüscher & Malenka, 2012). To do this, we evoked EPSCs at −70 mV and +40 mV to record AMPA receptor- and NMDA receptor-mediated currents, respectively. Recordings were made in the presence of 10μM SR95531 to block GABA_A_ receptors, thereby isolating excitation. Our results suggest that there was no significant difference in the AMPA/NMDA ratio across columns (vlPAG: 5.53±2.86; n=14, N=5; dmPAG: 7.22±4.73; n=14, N=3; paired t-test p = 0.26; **Figure 3A-C**), suggesting a similar density of AMPA/NMDA receptors across regions. To further characterize the potential receptor composition underlying these responses, we fit the EPSCs recorded at −70 mV and +40 mV with a double exponential. At −70 mV, the EPSC decay was significantly faster in the vlPAG (3.4±1.6 ms, n = 14, N = 5) than in the dmPAG (5.1±2.1 ms, n = 14, N = 3, paired t-test: p = 0.02; **Figure 3D**), which may suggest a difference in AMPA receptor subunit composition across columns. Of note, the kinetic profile of the synaptic responses recorded at +40 mV (reflecting a mix of AMPA and NMDA receptor components) were relatively fast in both the vlPAG (25.90±20.83 ms; n=14, N=5) and dmPAG (3.99±28.04 ms; n=14, N = 3) and were not significantly different (paired t-test p = 0.44, **Figure 3E**). The relatively fast time constant recorded at +40 mV suggests that the NMDA receptor subunit composition may be dominated by GluN2A-containing NMDA receptors, which have appreciably faster decay kinetics than other GluN2 subunits (Monyer *et al*., 1994; Vicini *et al*., 1998; Wyllie *et al*., 1998; Cull-Candy *et al*., 2001). Further, the faster kinetics likely resulted in underestimating the relative strength of NMDA receptor-mediated currents, as responses were measured 40 ms after stimulus onset.

**Figure 3.**
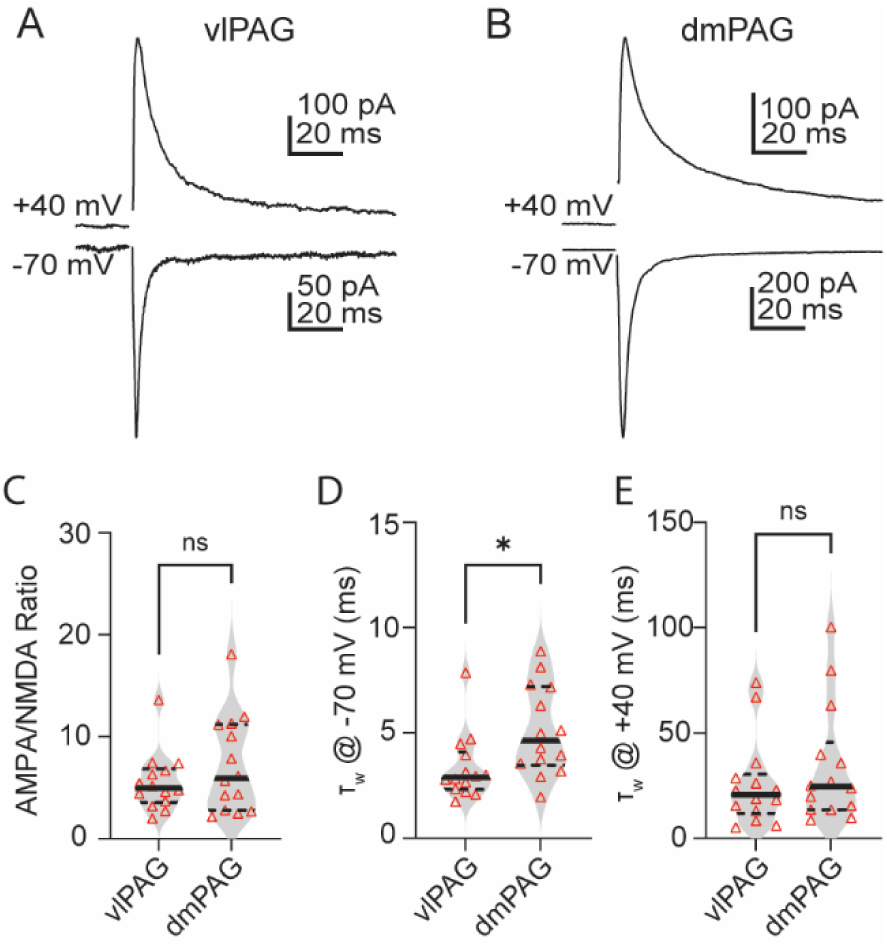
AMPA to NMDA ratio in the vl- and dmPAG. **(A, B)** Representative traces of evoked postsynaptic currents recorded at −70 mV and +40 mV from a single cell in the vlPAG (A) and dmPAG (B). **(C)** Violin plot of AMPA/NMDA ratio across vl- and dmPAG columns. **(D)** Violin plot of weighted decay time constants, τ, between vl- and dmPAG measured at −70 mV. **(E)** Comparison of weighted decay time constants, τ, between vl- and dmPAG measured at +40 mV.

### Stability of paired pulse ratio across interstimulus intervals

Another important feature of circuit-level information processing is the profile of presynaptic release probability, as synapses with high or low release probabilities can impose distinct temporal filters (i.e. low-pass or high-pass filters, respectively) on information transfer. One common, albeit crude, measure of presynaptic release probability is the paired pulse ratio, which assumes negligible contributions of postsynaptic mechanisms (i.e. receptor desensitization) to synaptic amplitudes on relatively short timescales. Therefore, to assess presynaptic release probability in the vl- and dmPAG, we examined the paired pulse ratio across six interstimulus intervals (10, 20, 50, 100, 250, and 500 ms). Interestingly, in both columns, we observed relatively stable paired pulse ratios across all interstimulus intervals tested, suggestive of frequency invariant synaptic coding strategies. More specifically, in the vlPAG, while we observed modest paired pulse depression (i.e. a paired pulse ratio significantly less than 1) at interstimulus intervals <250 ms (**Table 2**), the paired pulse ratio did not vary as a function of interstimulus interval (one-way ANOVA: p = 0.40; **Figure 4A, B**). A similar trend was observed for inhibitory paired pulse ratios in the vlPAG, which showed similar frequency invariance (one-way ANOVA: p = 0.0514; **Figure 4A, C**) but weaker overall depression (**Figure 4A, C).**

**Figure 4.**
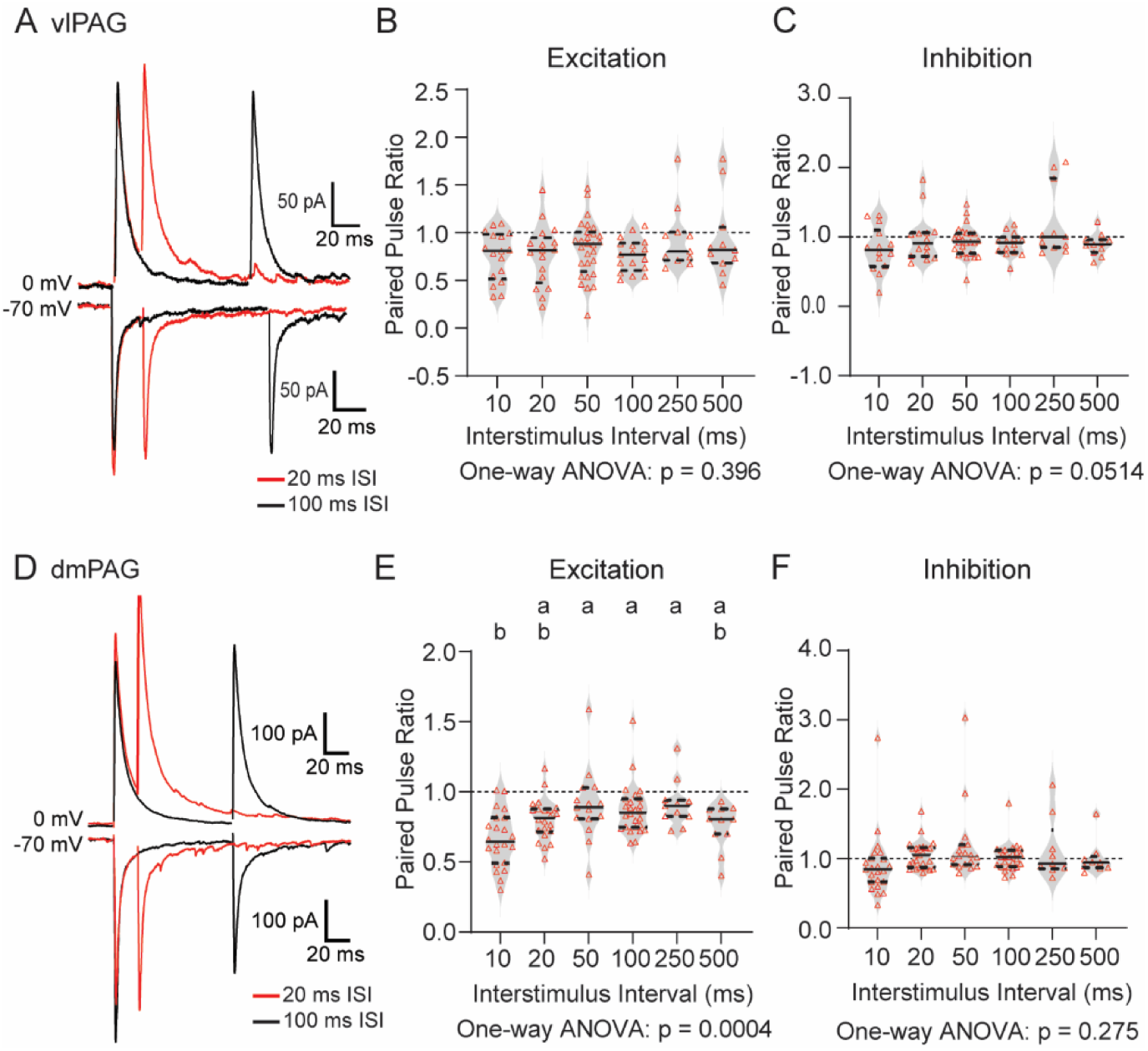
Evidence of frequency invariant coding in PAG columns. **(A)** Representative evoked postsynaptic currents showing excitatory and inhibitory paired pulse ratios in the vlPAG at 20 ms interstimulus intervals (red) and 100 ms interstimulus intervals (black). **(B, C)** Distribution of excitatory (B) and inhibitory (C) paired pulse ratio across interstimulus intervals (10 – 500 ms) in vlPAG. **(D)** Representative evoked postsynaptic currents showing excitatory and inhibitory paired pulse ratios in the dmPAG. **(E, F)** Distribution of excitatory (E) and inhibitory (F) paired pulse ratio across interstimulus intervals (10 – 500 ms) in dmPAG.

**Table 2:** Paired Pulse Ratios. Excitatory and inhibitory paired pulse ratios (mean±S.D.) across columns and interstimulus intervals. Level of significance indicates result of a one-sample t-test against a hypothetical mean of 1.0.

| Column | vIPAG |  |  |  | dmPAG |  |  |  |
| --- | --- | --- | --- | --- | --- | --- | --- | --- |
|  | eEPSC PPR |  | eIPSC PPR |  | eEPSC PPR |  | eIPSC (PPR) |  |
| ISI (ms) | Mean<br>(n) | p | Mean<br>(n) | p | Mean<br>(n) | p | Mean<br>(n) | p |
| 10 | 0.76±0.26<br>(17) | * | 0.82±0.33<br>(17) | ns | 0.66±0.20<br>(22) | **** | 0.91±0.47<br>(23) | ns |
| 20 | 0.77±0.32<br>(17) | ** | 0.98±0.35<br>(17) | ns | 0.81±0.15<br>(22) | **** | 1.05±0.20<br>(25) | ns |
| 50 | 0.83±0.30<br>(28) | ** | 0.94±0.23<br>(28) | ns | 0.90±0.26<br>(15) | ns | 1.18±0.53<br>(18) | ns |
| 100 | 0.76±0.18<br>(17) | **** | 0.90±0.17<br>(17) | * | 0.87±0.18<br>(26) | ** | 1.02±0.21<br>(25) | ns |
| 250 | 0.94±0.33<br>(11) | ns | 1.20±0.51<br>(11) | ns | 0.92±0.17<br>(11) | ns | 1.13±0.45<br>(9) | ns |
| 500 | 0.93±0.42<br>(11) | ns | 0.89±0.16<br>(11) | * | 0.76±0.16<br>(11) | *** | 1.00±0.25<br>(9) | ns |

Conversely, in the dmPAG, we observed significant variation in the excitatory paired pulse ratio as a function of the interstimulus interval. More specifically, we observed stronger paired pulse depression at the shortest interval tested (10 ms) which decreased at longer interstimulus intervals (one-way ANOVA: p = 0.0004; **Figure 4D, E**). Further, as in the vlPAG, neurons in the dmPAG showed modest paired pulse synaptic depression at each interval tested (**Table 2**). Unlike excitatory inputs, the inhibitory paired pulse ratio in the dmPAG was stable across all interstimulus intervals tested and did not significantly differ at any interval from a paired pulse ratio of 1 (one-way ANOVA: p = 0.27; **Figure 4D, F**). Taken together, these data suggest that the PAG, as a whole, prioritizes frequency invariant synaptic coding strategies with relatively little evidence of synaptic facilitation or depression across interstimulus intervals. This general coding strategy, however, does not apply to excitatory currents in the dmPAG, which show evidence of paired pulse depression at short interstimulus intervals.

### Excitatory synaptic responses to high frequency trains of stimulation

Although the paired pulse ratio is often used to assess synaptic facilitation or depression, frequency invariant coding strategies are better studied using high frequency stimulation, which can fatigue synapses and reveal distinct mechanisms underlying stable synaptic responses.

Therefore, we next used high frequency stimulation trains to probe the mechanisms of frequency invariant coding underlying excitatory synaptic transmission in the vl- and dmPAG. Consistent with frequency invariant synaptic coding, EPSCs recorded in the vlPAG generally remained stable across the stimulus train at each frequency tested (**Figure 5A-C**), suggesting relatively little contribution of synaptic depression and/or facilitation. To quantify the synaptic mechanisms contributing to such stability, we separately measured both the phasic and total current following each stimulus within the train (see methods). In the vlPAG, there was close alignment of the normalized phasic and total current across frequencies in each individual cell (**Figure 5D-F**). As a statistical measure, we compared the phasic and total current amplitude as a function of stimulus number, and observed no significant interaction between current type and stimulus number (**Figure 5G-I**; RM two-way ANOVA interaction term: 10 Hz: p = >0.99; 50 Hz: p = 0.30; 100 Hz: p = 0.42; n = 13, N = 6), indicating that any change in total current amplitude across stimuli was driven primarily by concomitant changes in the phasic current amplitude.

**Figure 5.**
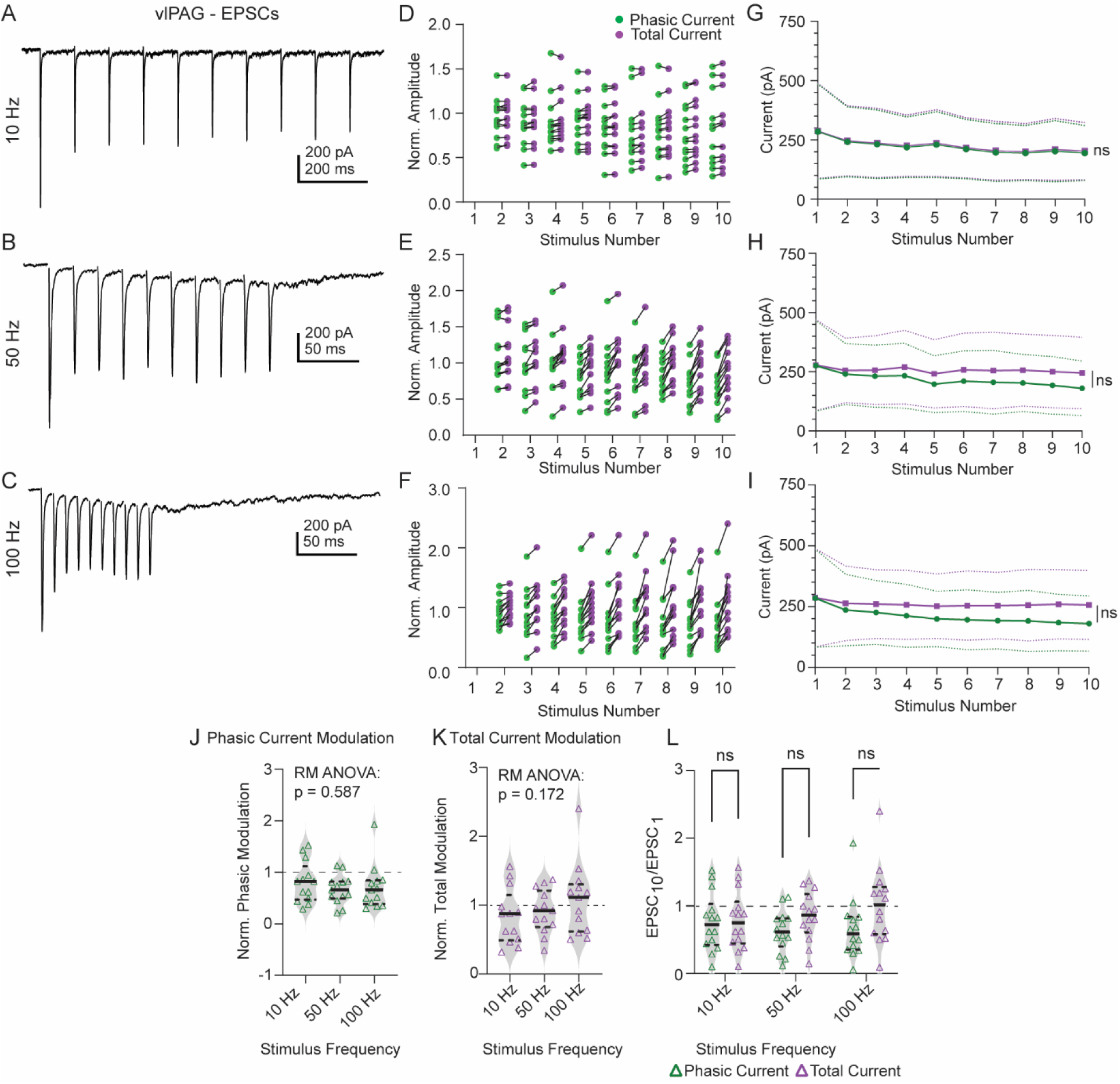
Excitatory synaptic dynamics across stimulation frequencies in the vlPAG. **(A-C)** Representative traces of evoked trains of excitatory postsynaptic currents in response to local electrical stimuli delivered at 10 Hz (A), 50 Hz (B), and 100 Hz (C) in the vlPAG. **(D-F)** Quantification of phasic (green) and total (purple) current amplitude at 10 Hz (D), 50 Hz (E), and 100 Hz (F), peak-normalized to first EPSC in train. Solid lines connect paired phasic and total current measurements from single cells. **(G-I)** Mean phasic (green) and total (purple) current amplitude plotted as a function of stimulus number at 10 Hz (G), 50 Hz (H), and 100 Hz (I). Dotted line represents 95% CI. **(J, K)** Distribution of peak-normalized phasic (J) and total (K) current amplitude across stimulation frequencies in the vlPAG. **(L)** Violin plot of phasic (green) and total (purple) current amplitude of last stimulus-evoked EPSC in train normalized to first EPSC in train across stimulation frequencies.

As an additional measure of the stability of the phasic and total current across the train, we compared phasic and total current modulation by normalizing the current recorded on stimulus 10 to the current recorded on stimulus 1. Consistent with frequency invariant synaptic coding, we observed no significant difference in either the normalized phasic (**Figure 5J**; 10 Hz: 0.81±0.40; 50 Hz: 0.66±0.28; 100 Hz: 0.71±0.43; RM one-way ANOVA: p = 0.59; n = 13, N = 6) or total (**Figure 5K**; 10 Hz: 0.84±0.40; 50 Hz: 0.91±0.32; 100 Hz: 1.1±0.52; RM one-way ANOVA: p = 0.45; n = 13, N = 6) current modulation across the stimulus train at any frequency tested. Additionally, at each stimulus frequency, the total current modulation across the stimulus train was not significantly different from 1 (one-sample t-test: 10 Hz: p = 0.17; 50 Hz: p = 0.32; 100 Hz: 0.60; n = 13, N = 6), supporting the conclusion that the total current amplitude did not significantly change across the stimulus train. Finally, we directly compared the phasic and total current modulation and observed no significant difference between currents at any stimulus frequency (**Figure 5L**; two-way ANOVA: main effect of frequency: p = 0.60; main effect of current type: p = 0.085; frequency x current interaction: 0.31). Taken together, these results suggest that excitatory synaptic transmission in the vlPAG utilizes frequency invariant coding strategies and further suggest that the invariant coding is achieved through stable phasic synaptic amplitudes across the frequencies tested.

We used a similar approach to examine frequency invariant synaptic coding in the dmPAG, which showed modest paired pulse depression at the shortest interstimulus interval tested. As in the vlPAG stimulation trains resulted in relatively stable *total* synaptic currents across all three frequencies tested (**Figure 6A-C**); however, the mechanism of frequency invariant coding appeared distinct to that observed in the vlPAG. More specifically, as stimulus frequency increased, we observed an increase in the degree of phasic current depression, which resulted in a mismatch between the normalized phasic and total current measured in each individual cell (**Figure 6D-F**). Such a mismatch between the normalized phasic and total current is suggestive of an underlying tonic current (or slow current) upon which the phasic currents are generated (**Figure 6A-C**). Consistent with this observation, when we plotted the raw phasic and total current amplitude as a function of stimulus number, at frequencies ≥50 Hz, there was a statistically significant deviation between phasic and total current (**Figure 6G-I**; RM two-way ANOVA interaction term: 10 Hz: p = 0.99; 50 Hz: p < 0.0001; 100 Hz: p < 0.0001; n = 15, N = 5). These data suggest that, within the dmPAG, the phasic current dynamics alone are insufficient to fully explain the relative stability of the total current amplitude across the train. Furthermore, while the phasic current showed increasing depression as a function of stimulus frequency (**Figure 6J**; 10 Hz: 0.69±0.14; 50 Hz: 0.49±0.29; 100 Hz: 30±0.18; RM one-way ANOVA: p < 0.0001; n = 15, N = 5), the total current remained stable across frequencies (**Figure 6K**; 10 Hz: 0.76±0.15; 50 Hz: 0.88±0.32; 100 Hz: 0.84±0.27; RM one-way ANOVA: p = 0.11; n = 15, N = 5). Despite the stability across frequencies in the dmPAG, the total current amplitude did show a modest degree of depression across the train (one-sample t-test: 10 Hz: p <0.0001; 50 Hz: p = 0.17; 100 Hz: 0.04; n = 15, N = 5). Finally, consistent with the observation of an additional slow current contributing to the total current amplitude in the dmPAG, the phasic and total current modulation significantly differed as a function of stimulus frequency (**Figure 6L**; two-way ANOVA: main effect of frequency: p = 0.0028; main effect of current type: p = 0.0006; frequency x current interaction: p <0.0001). Taken together, these results suggest that in the dmPAG, while frequency invariant coding strategies are employed, additional synaptic mechanisms are required to overcome the phasic current depression observed at higher stimulation frequencies.

**Figure 6.**
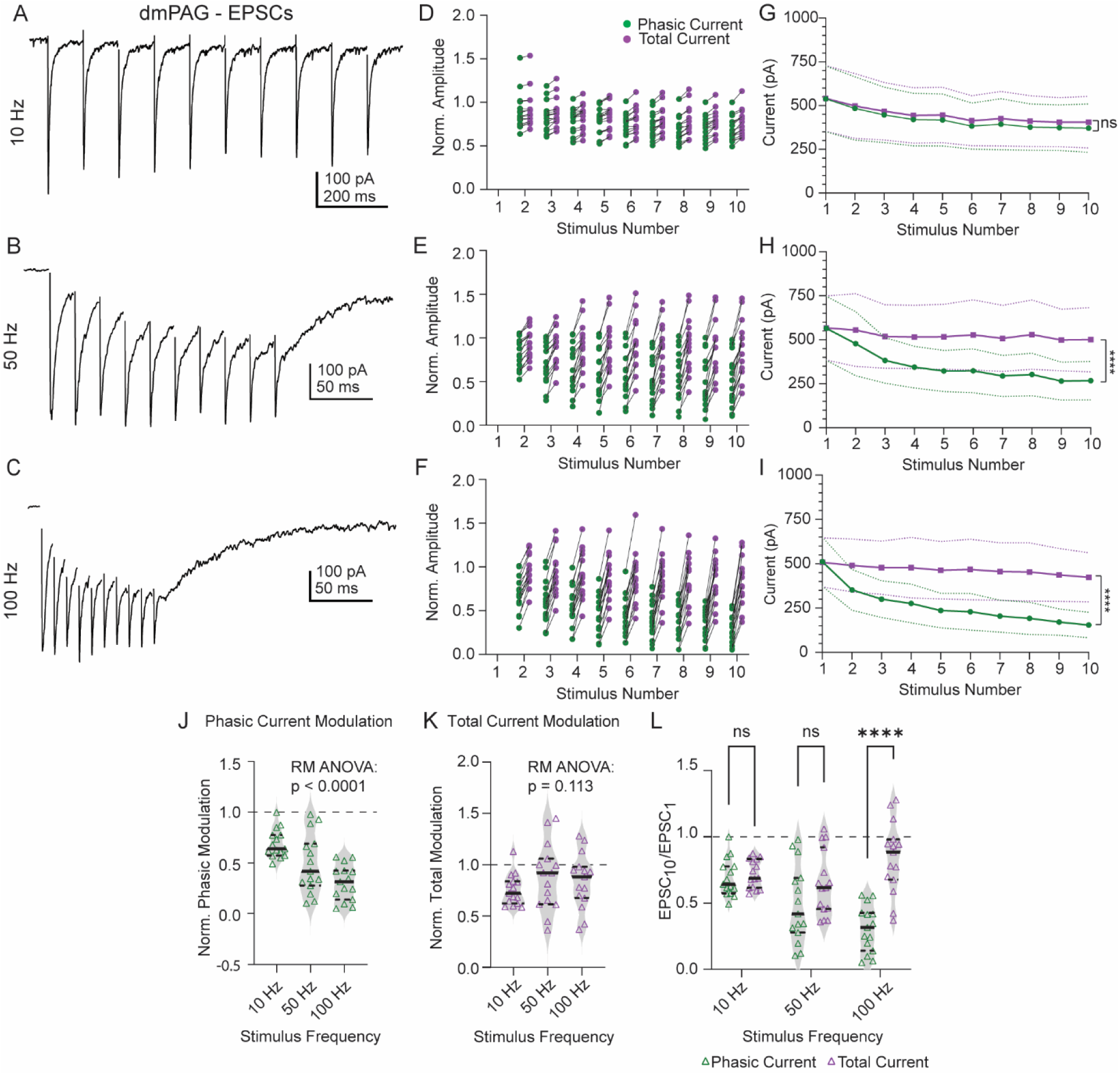
Excitatory synaptic dynamics across stimulation frequencies in the dmPAG. **(A-C)** Representative traces of evoked trains of excitatory postsynaptic currents in response to local electrical stimuli delivered at 10 Hz (A), 50 Hz (B), and 100 Hz (C) in the dmPAG. **(D-F)** Quantification of phasic (green) and total (purple) current amplitude at 10 Hz (D), 50 Hz (E), and 100 Hz (F), peak-normalized to first EPSC in train. Solid lines connect paired phasic and total current measurements from single cells. **(G-I)** Mean phasic (green) and total (purple) current amplitude plotted as a function of stimulus number at 10 Hz (G), 50 Hz (H), and 100 Hz (I). Dotted line represents 95% CI. **(J, K)** Distribution of peak-normalized phasic (J) and total (K) current amplitude across stimulation frequencies in the dmPAG. **(L)** Violin plot of phasic (green) and total (purple) current amplitude of last stimulus-evoked EPSC in train normalized to first EPSC in train across stimulation frequencies.

### Characterization and functional role of dmPAG tonic current

Our observations suggest that while both columns engage in frequency invariant synaptic coding strategies, the underlying mechanism necessary to achieve such invariance may differ across columns, specifically requiring a concomitant increase in slow current in the dmPAG to match phasic current depression. Therefore, we next sought to directly compare the tonic/slow current across columns. As expected, when we overlaid peak-normalized traces from the vl- and dmPAG (100 Hz), the tonic current amplitude both within and immediately following the stimulus train were significantly larger in the dmPAG (**Figure 7A**). To quantify these differences, we first measured the amplitude of the tonic current just prior to the last stimulus and normalized that value to the peak phasic current amplitude on stimulus 1. At frequencies ≥50 Hz, we observed a significantly increased tonic current buildup in the dmPAG compared to the vlPAG (**Figure 7B**; two-way ANOVA main effect of column: p = 0.0017; main effect of frequency p < 0.0001; Šídák multiple comparisons: 10 Hz: p = 0.94, 50 Hz: p = 0.046, 100 Hz: p = 0.031; vlPAG: n = 14, N = 6; dmPAG: n = 15, N = 5). These data suggest that during the stimulus train, there is a significantly larger increase in tonic current in the dmPAG. However, we also noted that slow current amplitude outlasted the stimulus train, which we quantified by measuring the total charge transfer in the 500 ms window immediately following 100 Hz stimulation. Consistent with enhanced tonic current in the dmPAG, the charge transfer in this window was significantly larger in the dmPAG (772.9±483.3 nC; n = 18, N = 5) than the vlPAG (132.6±107.6 nC, n = 14, N = 6; unpaired t-test: p <0.0001; **Figure 7C**). Finally, to assess how the tonic current developed throughout the stimulus train, we plotted the amplitude of the tonic current as a function of stimulus number. At all frequencies, we observed a significantly larger tonic current in the dmPAG compared to the vlPAG (**Figure 7D-F**; RM two-way ANOVA main effect of column: 10 Hz: 0.0006; 50 Hz: p = 0.0043; 100 Hz: p = 0.0004); additionally, there was a significant column x stimulus number interaction at each frequency tested (RM two-way ANOVA column x stimulus number interaction: 10 Hz p < 0.0001; 50 Hz: p < 0.0001; 100 Hz: p < 0.0001). These data confirm that the dmPAG has a significantly enhanced tonic current, which both counteracts the phasic current depression during the stimulus train (allowing for frequency invariant synaptic coding) and results in a prolonged window of slow excitation.

**Figure 7.**
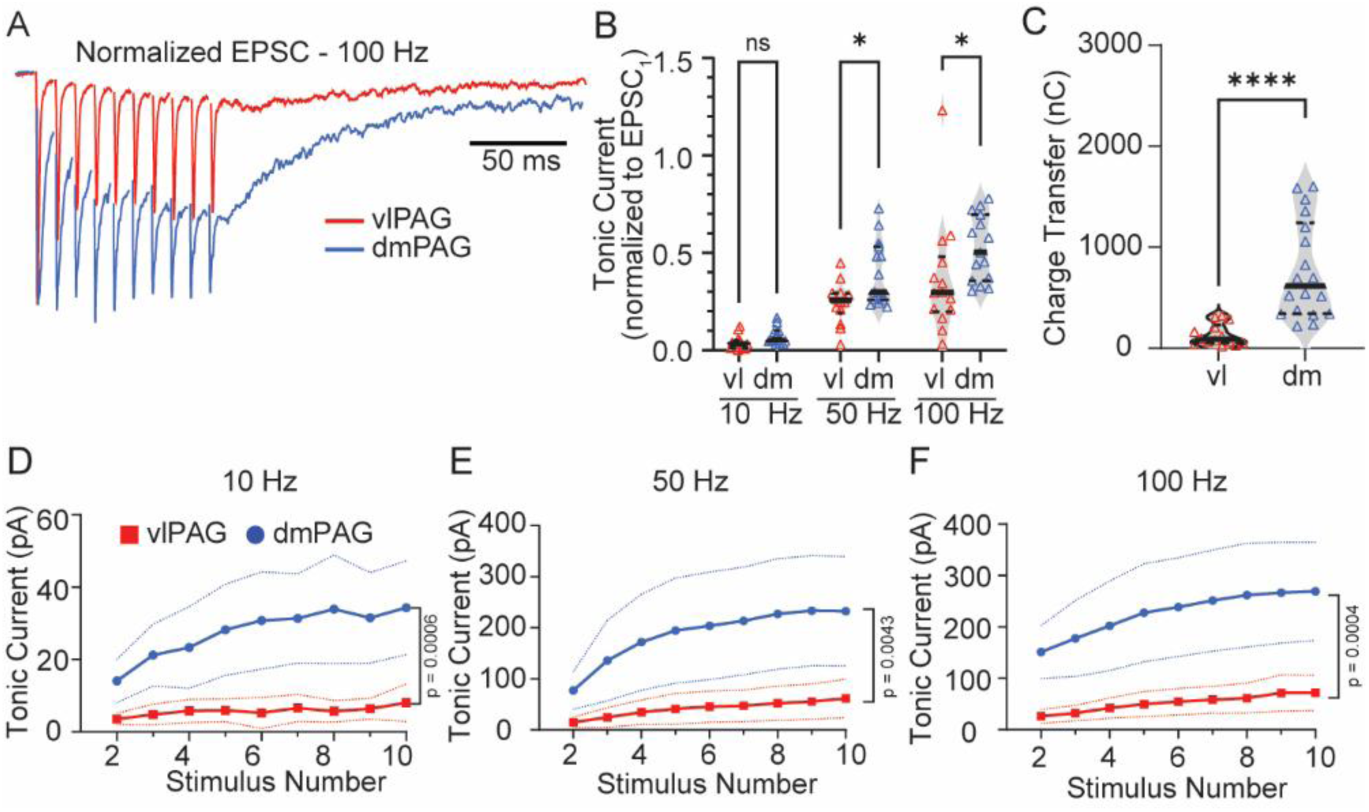
Comparison of tonic excitatory current between vl- and dmPAG columns in response to high frequency stimulation trains. **(A)** Representative traces of peak-normalized evoked trains of excitatory synaptic responses at 100 Hz in the vl- (red) and dmPAG (blue). **(B)** Violin plots comparing peak-normalized tonic current amplitude between columns across stimulation frequencies. **(C)** Violin plot of charge transfer measured over the 500 ms window following final stimulus in train in vl- and dmPAG. **(D-F)** Comparison of average tonic current amplitude in the vl- and dmPAG plotted as a function of stimulus number across frequencies. Dotted lines represent 95% CI.

To determine the functional impact of the prolonged synaptic excitation in the dmPAG, we recorded neurons in the dmPAG in current clamp configuration to determine whether the slow current resulted in prolonged firing. As expected, in the dmPAG, brief (5 stimuli) high frequency (100 Hz) stimulation resulted in prolonged spiking, which outlasted the duration of the stimulation epoch (**Figure 8A, B**). To quantify the change in firing rate, we calculated the firing rate immediately prior to (500 ms) and immediately following (1000 ms) high frequency stimulation, importantly excluding any action potentials generated during the high frequency stimulation itself. In all cells, we observed a significant increase in the firing rate after high frequency stimulation (**Figure 8C**; pre-stimulation: 17.18±17.05 Hz, post-stimulation: 31.30±23.06 Hz, paired t-test: p < 0.0001), which represented a 257.4±171.7% increase in firing rate (**Figure 8D**, one-sample t-test: p = 0.0016). These data suggests that the enhanced slow/tonic current observed in the dmPAG indeed results in a prolonged period of elevated firing rate following high frequency stimulation, which may contribute to the column-specific functions of the dmPAG in fear appraisal and behavior (Deng *et al*., 2016; Evans *et al*., 2018).

**Figure 8.**
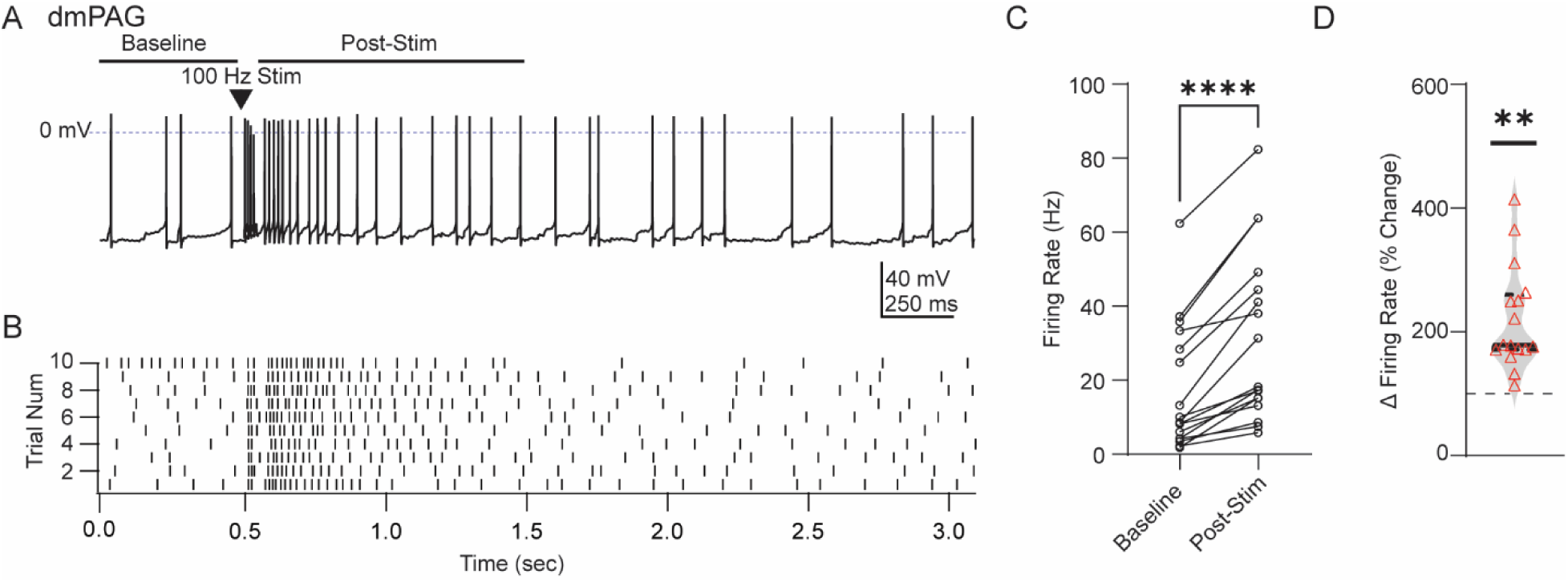
High frequency stimulation results in persistant increase in dmPAG firing rate. **(A)** Representative current clamp recording from a dmPAG neuron following brief (5 stimuli) high frequency (100 Hz) stimulation. **(B)** Raster plot illustrating persistant increase in firing rate in the ∼1 second period after high frequency stimulation. **(C)** Quantification of the firing rate before and after high frequency stimulation. **(D)** Violin plot of the percent change in firing rate following stimulation.

## Discussion

One of the current limitations in fully appreciating the diverse roles of the periaqueductal gray in fear processing is that the synaptic dynamics within and across columns is poorly understood. Therefore, our goal in this study was to directly examine and compare the intrinsic and synaptic properties of neurons across columns in the PAG. Our results identify circuit features that are conserved across PAG columns but also identify important differences in underlying synaptic dynamics which may contribute to distinct column function. Most strikingly, both vlPAG and dmPAG neurons engage in frequency invariant coding – demonstrating remarkably stable synaptic currents across multiple input frequencies. Despite this commonality, however, the underlying circuit mechanisms used to achieve such frequency invariance significantly differed between columns. In the vlPAG, the phasic current amplitude remained largely stable across stimuli at frequencies up to 100 Hz. Conversely, in the dmPAG, phasic current amplitudes showed significant depression at higher stimulation frequencies (50 and 100 Hz); however, this was matched by a buildup of tonic current across the stimulation train, resulting in stability of total current amplitudes across the train. These data provide important biophysical grounding for understanding column-specific PAG circuit function during threat processing.

### Comparison of intrinsic properties and basal synaptic transmission

Our results suggest that neurons in the dmPAG exhibited higher intrinsic excitability than neurons in the vlPAG. While there was no difference in the spontaneous firing rate across columns, neurons in the dmPAG exhibited a higher maximal firing in response to current injection and had action potentials with a shorter duration, which may contribute to the higher maximal firing rate. At the synaptic level, we observed gross similarities in the apparent local circuit arrangement across columns, with a consistent observation of larger spontaneous IPSCs, suggesting a larger unitary inhibitory amplitude in both columns. Interestingly, in the vlPAG, evoked inhibitory currents were significantly stronger than excitatory currents, suggesting a preferential extrinsic inhibitory input, potentially mediated by extrinsic vlPAG input from the central amygdala (Tovote *et al*., 2016). Together, these data suggest that while there may be similarities in the local circuit arrangement across columns, there are likely significant differences in how each column processes its diverse complement of extrinsic inputs (Beitz, 1982; Silva & McNaughton, 2019).

Our results also highlight important synaptic features which may constrain the extent of synaptic plasticity mechanisms within the PAG. NMDA-receptors are universally considered a key synaptic substrate for engaging long term synaptic potentiation and depression, owing to their role in calcium entry and subsequent biochemical cascades (Bear & Malenka, 1994; Malenka, 1994; Lüscher *et al*., 2000; Lüscher & Malenka, 2012). Ion flow through NMDA receptors requires both postsynaptic depolarization and glutamate binding, and generally lasts for hundreds of milliseconds due to the slow kinetic profile of NMDA receptors (Lester *et al*., 1990). Our synaptic data, however, suggest that the time course of NMDA receptor activation in the PAG is remarkably short, decaying with a time constant in the tens of milliseconds, likely reflecting NMDA receptors containing GluN2A subunits (Monyer *et al*., 1994; Wyllie *et al*., 1998; Cull-Candy *et al*., 2001). Such fast decay kinetics suggest that the time window for synaptic plasticity within the PAG may be relatively brief, as differences in the complement of synaptic receptors at the synapse can significantly impact plasticity profiles (Traynelis *et al*., 2010; Paoletti *et al*., 2013). Among glutamate receptors, precise tuning of kinetic profiles is primarily driven by the composition of the GluN2 subunit (Gielen *et al*., 2009; Yuan *et al*., 2009; Paoletti, 2011), and can influence the profile of synaptic plasticity (Evans *et al*., 2012).

### Frequency invariant synaptic efficacy and the underlying synaptic mechanisms

In many circuits, high frequency stimulation reveals a pattern of either facilitation or depression, which imposes circuit level transformations on information processing, through high- or low-pass synaptic filtering (Abbott *et al*., 1997; Abbott & Regehr, 2004). Our data, however, indicate relative stability of the paired pulse ratio across interstimulus intervals, which suggest relatively weak synaptic filtering of presynaptic input. Such frequency invariant synaptic coding is reminiscent of similar strategies imposed by other circuits including the cerebellum (Telgkamp *et al*., 2004; Bagnall *et al*., 2008; Turecek *et al*., 2016, 2017), where it is thought to facilitate high fidelity transmission across the circuit. As such, the relative frequency invariance observed here may be a synaptic mechanism to ensure that the PAG faithfully transmits and encodes presynaptic input, a feature consistent with a role in driving rapid behavioral responses to threats within the environment. Such a circuit feature would, in theory, require relatively minimal information processing within the PAG to ensure rapid and reliable behavioral execution. Despite this interpretation, it is important to note that our experiments utilized non-specific electrical stimulation, therefore obscuring the identity of the presynaptic neuron. Therefore, it is possible that the frequency invariance observed here reflects an averaging of facilitating and/or depressing synapses from distinct presynaptic sources. Future work is needed to specifically and systematically examine the synaptic dynamics of presynaptic inputs to the vl- and dmPAG from different presynaptic regions.

Despite the frequency invariance observed in both columns, the underlying synaptic mechanisms necessary to achieve this invariance significantly differed across columns. In the vlPAG, phasic currents were remarkably stable, with little deviation between phasic and total currents across stimulation trains. This stability suggests that frequency invariance in the vlPAG may be predominantly driven by an intermediate and relatively stable presynaptic release probability. In contrast, the dmPAG exhibited phasic current depression during high frequency stimulation trains (≥50 Hz), which was offset by a corresponding increase in tonic current. This tonic current persisted beyond the stimulation train and may be mediated by numerous mechanisms such as gap junctional coupling (Christie *et al*., 2005; Maher *et al*., 2009; Vaaga & Westbrook, 2016), recurrent excitatory networks (Molnár *et al*., 2008; Hemberger *et al*., 2019; Oberle *et al*., 2023), asynchronous release (Turecek *et al*., 2017), or slower metabotropic signaling. Functionally, the slow tonic current observed in the dmPAG aligns with previously described synaptic thresholding mechanisms within the dmPAG, selectively mediated by inputs from the superior colliculus (Evans *et al*., 2018). In fact, while our results were elicited by stimulation in the dorsal PAG, suggesting that a recurrent network may exist within the dmPAG itself, we cannot rule out the possibility that our stimulation antidromically activated the superior colliculus which may be preserved in our slice preparation and therefore contribute to the slow current recorded in the dmPAG.

### Relation to PAG columnar function

Classical models of PAG function ascribe distinct behavioral responses to threat to differential activation of specific columns within the PAG. More specifically, activation of the dorsal PAG is thought to coordinate active defensive strategies, such as flight, which are preferentially engaged by innate threats (Carrive, 1993; Bandler & Shipley, 1994; Vianna *et al*., 2001b, 2001a) whereas activation of the vlPAG is necessary to support freezing, specifically during fear conditioning paradigms (LeDoux *et al*., 1988; Kim *et al*., 1993; Carrive *et al*., 1997). However, more recent work using *in vivo* circuit manipulation and recording techniques during threat processing has revealed that this model of PAG function is likely too simplistic, as neurons in both columns process more complex features of threat.

For example, in the vlPAG, optogenetic stimulation of a molecularly defined (Chx10 positive) population of glutamatergic neurons results in robust freezing (Tovote *et al*., 2016; Vaaga *et al*., 2020) and in vivo recordings suggest that neurons in the vlPAG correlate with muscle activity during freezing in the period of early extinction after fear conditioning (Watson *et al*., 2016). However, more complex behavioral paradigms have also revealed that neurons in the vlPAG encode more abstract features of threat such as threat probability (Wright & McDannald, 2019) or a positive prediction error necessary for updating fear responses (Johansen *et al*., 2010; Roy *et al*., 2014; Groessl *et al*., 2018; Walker *et al*., 2019). It is worth noting that these two functions in fear processing need not be mutually exclusive, but it does suggest that the vlPAG is engaged in more complex features of threat processing, potentially in addition to its role in driving freezing behavioral output.

Similar complexities have also been observed in the dmPAG, which suggest that, rather than simply encoding flight behavior, neurons in dmPAG are involved in escape initiation, likely through interactions with locomotor centers including the mesencephalic locomotor region (Evans *et al*., 2018; Ferreira-Pinto *et al*., 2018). Such escape initiation, in fact, is thought to be mediated by a synaptic thresholding mechanism between the superior colliculus and the dmPAG that closely resembles our prolonged synaptic excitation preferentially observed in the dmPAG (Evans *et al*., 2018). Furthermore, neural recordings in the dorsal PAG reveal two populations of neurons: assessment cells that ramp their activity during threat assessment and flight cells whose activity was directly correlated with flight behavior (Deng *et al*., 2016). Further, molecularly distinct neurons in the lateral PAG, which express CCK, may directly encode flight behavior (La-Vu *et al*., 2022).

These emerging findings across columns highlight the need for a more nuanced understanding of PAG function that accounts for molecularly defined cell populations as well as the complexity in both the functional role of the PAG in threat processing and the underlying neural circuit dynamics. Overall, our findings provide important biophysical insights into the intrinsic and synaptic properties of neurons across columns of the PAG and highlight both conserved elements and distinct features of PAG column circuit dynamics that may relate to their shared and unique roles in driving behavioral action and engaging more nuanced threat processing.

## Author contributions

Ana Valeria Castro Romero: investigation, methodology, formal analysis, writing – original draft, writing – review and editing, visualization; Kate Tyner: investigation, methodology, formal analysis, writing – original draft, writing – review and editing, visualization; Jordan Carroll: investigation, formal analysis, writing – original draft, writing – review and editing; Christopher Vaaga: conceptualization, methodology, formal analysis, investigation, writing – original draft, writing – review and editing, supervision, funding acquisition.

## Acknowledgments

We thank members of the Vaaga lab for helpful discussion and feedback on the project.

## Declarations

The authors have nothing to declare.

